# Planktonic interference and biofilm alliance between aggregation substance and endocarditis and biofilm associated pili in *Enterococcus faecalis*

**DOI:** 10.1101/348029

**Authors:** Irina Afonina, Xin Ni Lim, Rosalind Tan, Kimberly A. Kline

## Abstract

Like many bacteria, *Enterococcus faecalis* encodes a number of adhesins involved in colonization or infection of different niches. Two well-studied *E. faecalis* adhesins, aggregation substance (AS) and endocarditis and biofilm-associated pili (Ebp), both contribute to biofilm formation on abiotic surfaces and in endocarditis, suggesting they may be expressed at the same time. Because different regulatory pathways have been reported for AS and Ebp, here we examined if they are co-expressed on the same cells and the functional impact of co-expression on individual cells and within a population. We found that while Ebp are only expressed on a subset of cells, when Ebp and AS are expressed on the same cells, pili interfere with AS-mediated clumping and impede AS-mediated conjugative plasmid transfer during planktonic growth. However, when the population density increases, horizontal gene transfer rates normalize and are no longer affected by pilus expression. Instead, at higher cell densities during biofilm formation, Ebp and AS differentially contribute to biofilm development and structure, synergizing to promote maximal biofilm formation.

## IMPORTANCE

Most bacteria express multiple various adhesins that contribute to surface attachment and colonization. However, the network and relationships between multiple adhesins of a single bacterial species are less-well understood. Here we examined two well-characterized adhesins in *Enterococcus faecalis*, aggregation substance and endocarditis and biofilm-associated pili, and found that they exhibit distinct functional contributions depending on the growth stage of the bacterial community. Pili interfere with aggregation substance-mediated clumping and plasmid transfer under planktonic conditions, whereas both adhesins structurally complement each other during the biofilm development. Together this study advances our understanding of how *E. faecalis*, a ubiquitous member of the human gut microbiome and an opportunistic pathogen, uses multiple surface structures to evolve and thrive.

## INTRODUCTION

*Enterococcus faecalis* is a Gram-positive opportunistic pathogen that causes a variety of infections such as endocarditis, bacteremia, urinary tract infections (UTI), and catheter associated urinary tract infections (CAUTI) (1–3). Most infections start with bacterial adhesion to a biotic or abiotic surface, and *E. faecalis* encodes multiple adhesins that facilitate attachment to and colonization of different niches within the host. Sortase enzymes are conserved within Gram-positive bacteria and catalyze the covalent attachment of many adhesins to the cell wall. Sortase substrates can be predicted based on the presence of a conserved sortase A recognition motif, LPxTG (leucine, proline, x = any amino acid, threonine and glycine), within a canonical cell wall sorting signal (4). *E. faecalis* strain V583 encodes 41 predicted SrtA substrates (5). Of these putative substrates, 17 are predicted to be MSCRAMM (microbial surface component recognizing adhesive matrix molecules), although only a few, including endocarditis* and biofilm-associated pili (Ebp) and aggregation substance (AS), have been characterized in detail (2, 6–8).

Ebp is composed of 3 subunits – EbpA, EbpB and EbpC, where EbpC is the major pilus subunit with EbpB on the base and EbpA on the tip of the pilus (2, 9). The three subunits are cotranscribed at the *ebpABC* locus and are positively regulated by the transcriptional regulator EbpR, which is encoded upstream of *ebpABC* (10). Polymerized Ebp exist as high molecular weight polymers (>200 kDa) and the length of the pilus may reach 10 μm (5). Ebp are only expressed on a subset of cells in the population, suggesting they may be phase variable, and pilus expression can be induced by exposure to serum, glucose, or bicarbonate (2, 11–13). EbpA mediates attachment to host fibrinogen and collagen, and contributes to UTI, CAUTI, and endocarditis (2, 3, 13). Mutations in the EbpA tip adhesin prevent Ebp-associated biofilm formation *in vitro* as well as to CAUTI in mice (2, 3, 14).

AS is 137 kDa protein encoded by *prgB* on the pheromone-responsive pCF10 plasmid (15). In the absence of the cCF10 pheromone, expression of AS and most of the pCF10 plasmid-encoded genes are inhibited by the small plasmid-encoded and constitutively expressed peptide iCF10 (16). Alternatively, expression of AS can be induced by albumin-lipid complexes in the bloodstream that sequester or degrade iCF10, resulting in the activation of autocrine pheromone signaling (17). Scanning electron microscopy experiments demonstrated that AS is also only expressed on a subset of cells in a population, even at saturating concentrations of the cCF10 pheromone (18). When expressed, AS contributes to biofilm formation, cellular aggregation required for conjugative plasmid transfer, and increased virulence in endocarditis models (1, 19–21). In addition, AS facilitates adherence of *E. faecalis* to renal tubular cells, intestinal epithelial cells, as well as binding to and survival in neutrophils (22).

Most bacteria encode multiple adhesins; however, they are not always expressed at the same time, and this differential expression can arise via cross-regulation. For example, *E. coli* Pap pili and Type I fimbriae are cross-regulated, as are flagella and type IV pili in *P. aeruginosa* (23–25). Despite an increasing number of characterized and predicted adhesins in *E. faecalis*, we do not know whether or how adhesin expression is coordinated within a population. In this study, we used the well-characterized *E. faecalis* adhesins Ebp and AS to test the hypothesis that Ebp and AS are differentially expressed on different cellular subsets, which may give rise to partitioned adhesive functions within a population. Contrary to our hypothesis, we detected no transcriptional cross-regulation between the two adhesins, and instead observed rapid expression of AS in pheromone-induced cultures on nearly all cells, with Ebp co-expression on a subset of those cells. Simultaneous expression of Ebp and AS on the same cells prevented AS-mediated clumping required for conjugative plasmid transfer and, consequently, reduced horizontal gene transfer (HGT). Within biofilms, we demonstrate distinct functional contributions of Ebp and AS to biofilm development and structure, working synergistically to promote biofilm formation.

## RESULTS

### AS and Ebp are co-expressed on the same cells after pheromone induction

Previous studies reported that neither AS nor Ebp are expressed on all cells within a population, but that both adhesins contribute to biofilm formation (2, 19, 20, 26). We therefore hypothesized that expression of the two adhesins may be coordinated within a population such that different population subsets express different adhesin repertoires for optimal colonization, virulence, or biofilm architecture development. To address this hypothesis, we first quantified the expression of AS and Ebp by diluting overnight cultures of *E. faecalis* strain OG1RF (a rifampicin and fusidic acid resistant derivative of OG1) into fresh media containing the cCF10 pheromone (0.12 ng/ml).

After 30 min of pheromone exposure, 82% of the cells expressed AS on the cell surface, and this number increased to 95% by 90 min of pheromone exposure (**Figure 1A**). The fraction of AS-expressing cells was higher than we expected based on earlier reports in the same strain in which representative scanning electron micrographs (SEM) of exponentially grown, pheromone-induced bacteria showed only ^~^75% of cells expressed AS (18), which may be due to the increased sensitivity of AS detection by immunofluorescent microscopy (IFM) compared to SEM. After 18 hours of growth in the presence of cCF10, the percentage of AS-expressing cells in the population dropped significantly to 4% (**Figure 1A**). Since AS transcription peaks between 30-60 min after addition of pheromone and returns to uninduced levels by the end of the second hour, the drop in the number of AS-expressing cell is likely due to accumulation of inhibitor iCF10, along with division and dilution of AS-expressing cells coupled with replacement by new AS uninduced daughter cells (17, 27). By contrast, Ebp were expressed only on 20-40% of cells, and Ebp expression was independent of cCF10 exposure or AS expression (**Figure 1B**). Therefore, while we originally hypothesized that Ebp and AS expression might be restricted to distinct population subsets, since pheromone induction resulted in AS expression on the majority of cells and did not affect variable Ebp expression, we revised our hypothesis and predicted that AS and Ebp would be co-expressed together on a subset of cells. Indeed, when we performed co-immunostaining using AS and EbpC antiserum on pheromone induced cells, we observed that the two adhesins were displayed on the same cells and co-localized at the same hemispherical areas of the cell (**Figure 1C**).

**Figure 1.**
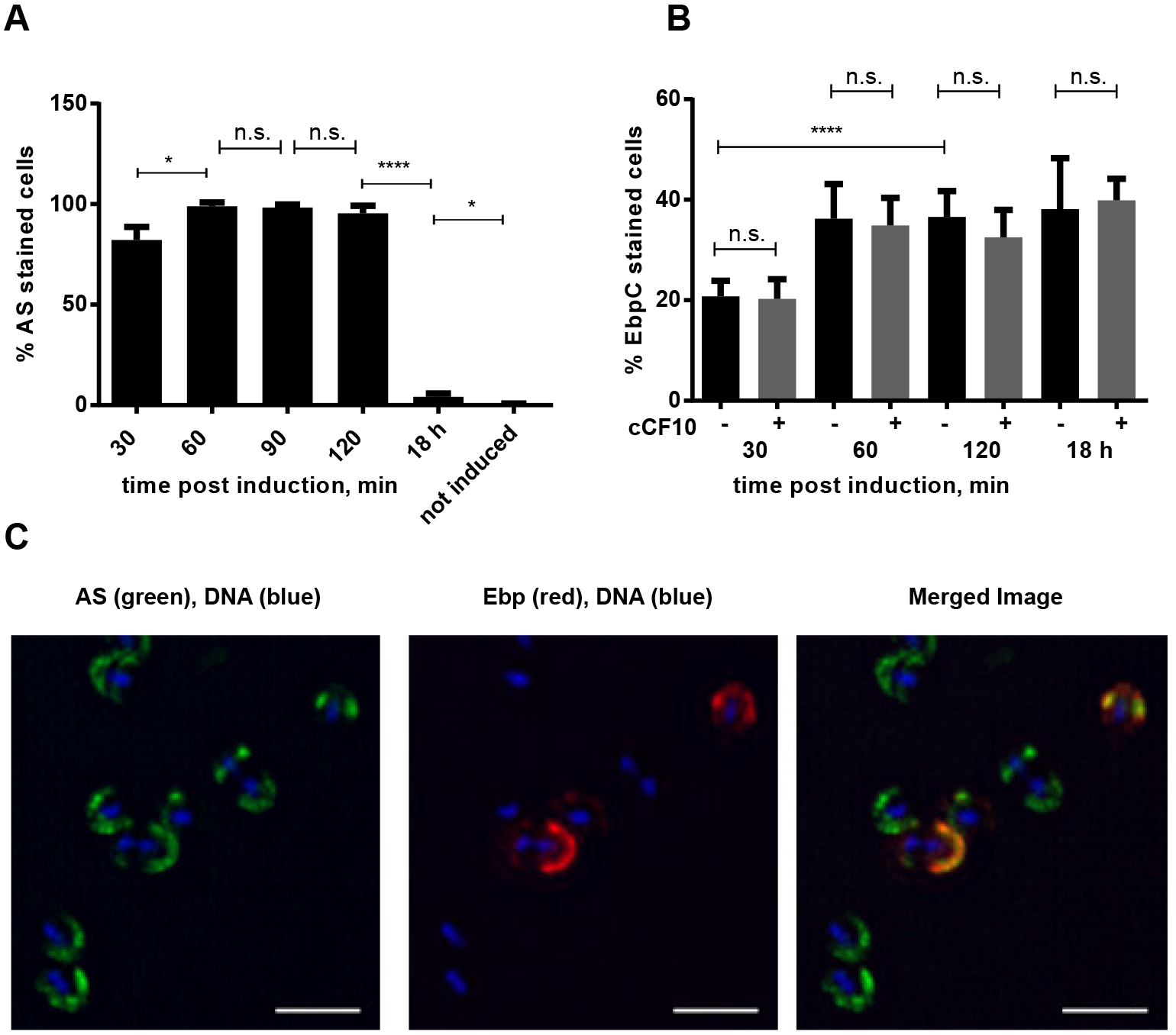
AS and Ebp are co-expressed on the same cells after pheromone induction. **A**. IFM was performed with AS antiserum and the % of AS+ cells within the population was quantified. **B**. IFM with EbpC antiserum was performed on cCF10 pheromone uninduced (black bars) and induced (grey bars) OG1RF pCF10 at the indicated time points and the % of EbpC+ cells within the population was quantified. **C**. IFM on pheromone induced cells with Ebp antiserum (red), AS (green), DNA (blue). Scale bar 1 μm. For **A** and **B**, the mean values are shown from 3 independent experiments in which at least 300 of cells were counted. Error bars represent the standard deviation. Statistical analysis was performed by the unpaired t-test using GraphPad. *p<0.05, ****p<0.0001, n.s.: p>0.05.

### Ebp interfere with AS-mediated clumping

AS was originally described for its association with cellular aggregation (28). While we did not observe a difference in Ebp expression between AS-expressing and AS-non-expressing populations (**Figure 1B**), we noticed that not all cells in cCF10-induced cultures aggregated, but instead separated into clumped cells that settled at the bottom of the tube as a pellet and cells that remained in suspension. Separation became obvious 1.5 - 2 hours after addition of cCF10 to the cultures (**Figure 2A**).

**Figure 2.**
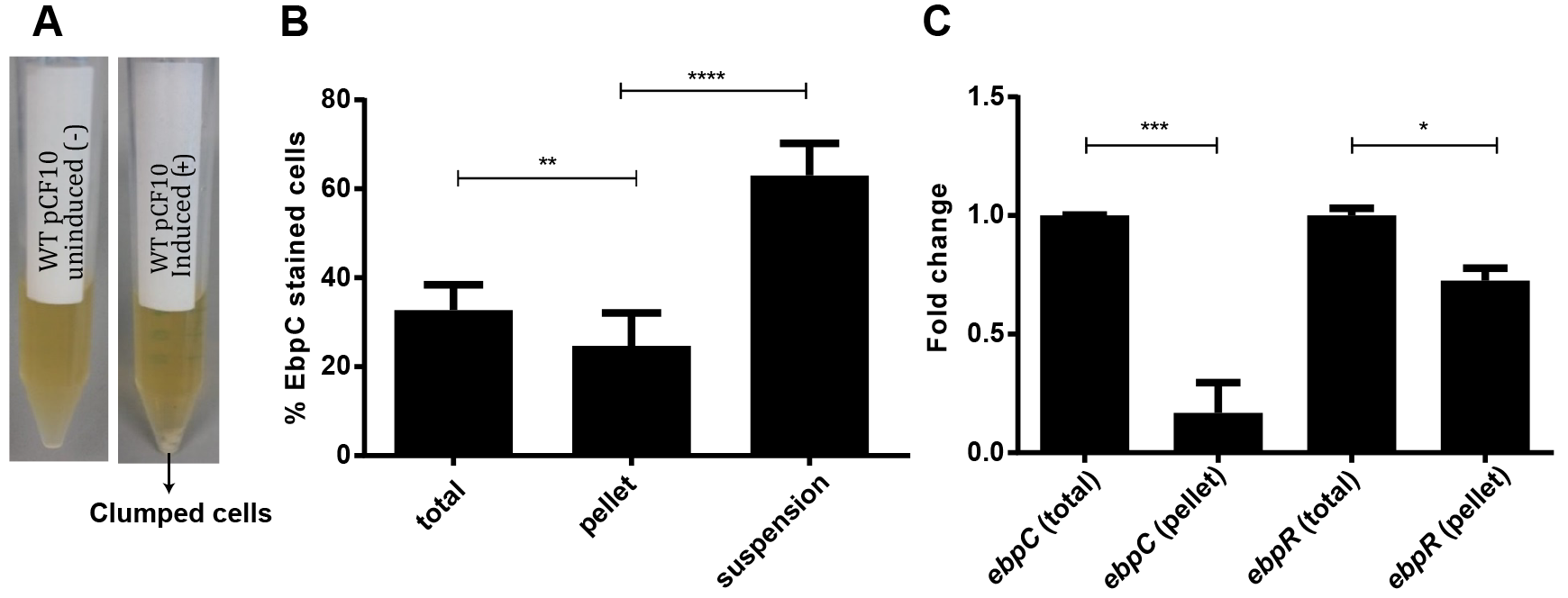
Ebp interfere with AS-mediated clumping. **A.** Representative image of OG1RF pCF10 uninduced (left) and OG1RF pCF10 induced (right) with pheromone cCF10 (0.12 ng/ml), 2 hours after induction. **B.** IFM was performed 2 hours post cCF10-induction with EbpC antiserum on the total population, suspension fraction (top 2 ml), and pellet fraction (clumped cells) and the % of EbpC+ cells was quantified. **C**. RT-qPCR on RNA isolated from pheromone induced cells two hours post cCF10 induction. Fold change indicates the change in *ebpC* and *ebpR* transcription in the pellet cells, compared to control (total). Mean values are shown from 3 independent experiments. Error bars represent the standard deviation. Each experiment was performed in triplicate using the ΔΔCt method to assess expression changes with *gyrA* as the reference housekeeping gene. Statistical analysis was performed by unpaired t-test using GraphPad.* p<0.05, ** p< 0.01, *** p<0.001; **** p<0.0001, n.s.: p>0.05.

Since Ebp in *E. faecalis* also contributes to cellular aggregation (29), we hypothesized that the clumping we observed in pheromone-induced cultures might be both AS- and Ebp-dependent, where pilus expressing cells might facilitate AS-mediated clumping. To test this, we manually separated the clumped pellet from the suspended cells, stained them with Ebp antiserum, and performed IFM. Contrary to our hypothesis, we observed fewer Ebp-expressing cells in the clumped pellet and more in the suspension when compared to the total (mixed) induced culture where pellets and suspended cells were not separated (**Figure 2B**).

We next performed RT-qPCR on the pellet cells and confirmed that *ebpC* transcript levels were lower in the pellet cells compared to the total population, which correlated with a similar transcript reduction of *ebpR*, a positive transcriptional regulator of *ebp* genes (**Figure 2C**). Given that Ebp expression levels within the total population was unchanged upon cCF10 pheromone exposure (**Figure 1B**), we speculated that Ebp-expressing cells may physically segregate to the suspension and be excluded from the aggregated pellet and we hypothesized that pilus-mediated steric hindrance could interfere with AS-mediated clumping.

### Ebp impede AS-facilitated horizontal gene transfer

Since AS-mediated clumping facilitates conjugative transfer of the pCF10 plasmid to recipient cells (20), we hypothesized that non-clumping suspension cells, which express both AS and Ebp, would display a reduced ability to undergo AS-mediated plasmid transfer compared to the pellet cells. To test this, we quantified conjugation frequency using suspension (Ebp^hi^ AS+) or pellet (Ebp^lo^ AS+) cells using OG1SS pCF10 as a donor (a streptomycin and spectinomycin resistant derivative of OG1 carrying pCF10 which encodes tetracycline resistance). After two hours of pheromone exposure, we collected the top 2 ml of each culture, containing suspension cells and the bottom 1 ml of clumped cells containing the pellet, out of an overall 5 ml culture. We normalized the bacterial cell numbers and mixed them with OG1RFΔ*ebpABCsrtC* recipient cells that lack pCF10 (rifampicin and fusidic acid resistant). We used Ebp deficient recipient cells to avoid pilus-mediated interference when incubated with the donor cells. Consistent with our hypothesis, 30 min of co-incubation of Ebp^lo^ AS+ pellet donor cells and Ebp null recipient cells yielded approximately 10 times more transconjugants compared to the Ebp^hi^ AS+ suspension donor cell mixture (**Figure 3A**).

**Figure 3.**
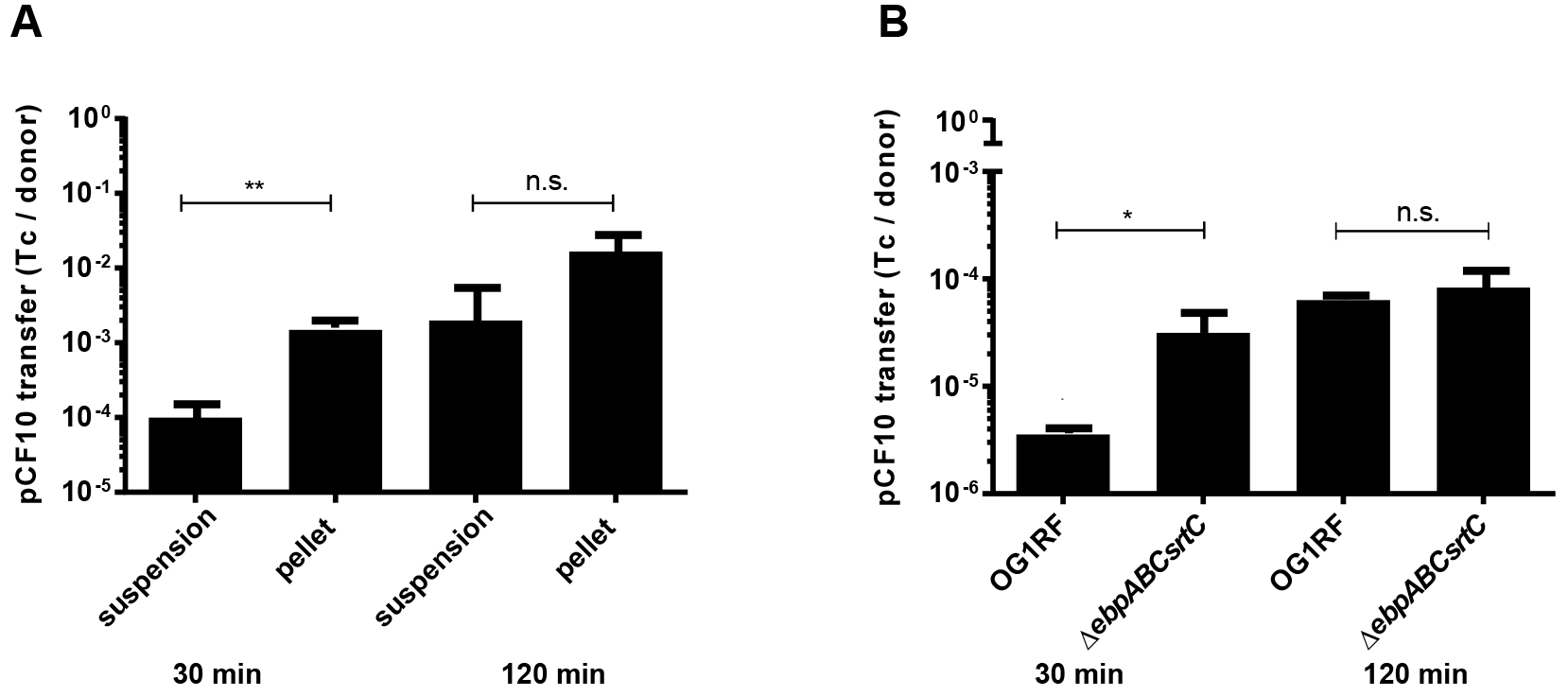
Ebp impede AS-mediated HGT. **A.** HGT rates from suspension and pellet cells of donor OG1SS pCF10 (str and tet resistant, rif sensitive) 2 hours post pheromone induction to plasmid free, Ebp null recipient OGlRFΔ*ebpABCsrtC* (rif resistant, str and tet sensitive). **B**. HGT rates from OG1RF pCF10 and Δ*ebpABCsrtC* pCF10 (both rif and tet resistant, str sensitive) donor cells to plasmid free recipient OGlXΔ*srtC* (str resistant, rif and tet sensitive). HGT rate is expressed as the number of transconjugants (Tc) per donor cell 30 and 120 min post mating. Error bars represent the standard deviation from 3 independent experiments. Statistical analysis was performed by the unpaired *t-test* using GraphPad. * p<0.05, ** p< 0.01, n.s.: p>0.05.

To further test the hypothesis that pili interfere with AS-mediated HGT, we used either OG1RF pCF10 or an OG1RF Δ*ebpABC* pCF10 (ebp null) as the donor strain and compared pCF10 conjugation rates to a OG1XΔ*srtC* recipient (streptomycin resistant) that fails to polymerize pili (**Figure 3B**). After 30 min of co-incubation, the Δ*ebpABC* strain was approximately 10-fold more efficient in plasmid transfer to OG1XΔ*srtC* as compared to OG1RF. In another study, similar conjugation experiments performed after 2 hours of co-culture revealed no differences in HGT for Ebp null donor strains compared to OG1RF (20). To address this, we extended our assay from 30 min to 2 hours and also observed that HGT rates equalized (**Figure 3A, B**). This delayed equalization of recovered transconjugants may be due to a second round of transfer of the plasmid from new donors to recipient cells, a switch to “off” piliation of originally “on” piliation donor cells, and/or enhanced cellular density that facilitates cell-to-cell contact and HGT.

### AS-clumped cells mediate microcolony formation and biofilm development

Both AS and Ebp contribute to biofilm formation, and pheromone induction of OG1RF pCF10 gives rise to thicker biofilms compared to OG1RF without the plasmid (20). Since we observed that AS-expressing cells clump after 2 hours of pheromone exposure, and those clumps largely exclude Ebp-expressing cells, we hypothesized that the initial attachment stage of biofilm formation by OG1RF pCF10 cells is mediated by AS clumps to form microcolonies, and that Ebp may be important at later stages to enhance biofilm maturation. To test this hypothesis, we quantified the number of Ebp-expressing cells attached and incorporated into early biofilms, after 2 hours of pheromone exposure, comparing OG1RF and OG1RF pCF10. In agreement with our hypothesis, we observed that the number of Ebp-expressing cells was lower in OG1RF pCF10 (12%) compare to OG1RF (22%), suggesting that if AS is present, AS aggregates will promote biofilm formation to the exclusion of Ebp-expressing cells (**Figure 4A**). To further explore the contributions of Ebp and AS, together and individually, to biofilm development we assayed biofilm formation by OG1RF (Ebp+ AS−), OG1RF pCF10 (Ebp+, AS+), OG1RFΔ*ebpABCsrtC* (Ebp−, AS−) and OG1RFΔ*ebpABCsrtC* pCF10 (Ebp-, AS+) (**Figure 4B**). Consistent with previous reports (2), we observed reduced biofilm formation by OG1RFΔ*ebpABCsrtC* as compared to OG1RF, and increased biomass of the induced OG1RF pCF10 that expresses both AS and Ebp as compared to OG1RF (20) (**Figure 4B**). Interestingly, we observed similar biofilm biomass for OG1RF and induced OG1RFΔ*ebpABCsrtC* pCF10, suggesting that in the absence of Ebp, AS alone is sufficient to revert the biomass of the *ebpABC* null strain to wild type levels (**Figure 4B**). To determine how Ebp and AS contribute to biofilm structure and organization, we grew biofilms for 24 hours in the presence of cCF10, stained the DNA of all of the cells with Hoechst, and performed confocal laser scanning microscopy. We observed distinct differences in the biofilm structure and thickness between the strains that express one, both, or neither of the adhesins (**Figure 5A, B**). In the absence of AS, we observed OG1RF to form uniform and tightly packed biofilms, while AS-expressing OG1RF pCF10 cells formed additional 3-dimensional microcolonies suggesting that AS promotes the formation of structured biofilm (**Figure 5A, B**). Moreover, while we observed uniform, tightly packed biofilms in Ebp-expressing OG1RF and OG1RF pCF10, the Δ*ebpABCsrtC* and Δ*ebpABCsrtC* pCF10 Ebp-null strains appeared more sparsely packed (**Figure 5A, B**), suggesting that Ebp are important for bacterial convergence. Furthermore, we observed that AS drives microcolony formation through cellular clumping, since we noticed elevated microcolony clusters only for pCF10-containing, AS-expressing strains (**Figure 5B**). Finally, Ebp contributed to the development of a thicker biofilms in OG1RF and AS-expressing OG1RF pCF10 when compared to Ebp-null counterparts (**Figure 5B**). We therefore conclude that despite accumulating similar overall biofilm biomass (**Figure 4B**), OG1RF and OG1RFΔ*ebpsrtC* pCF10 strains that display opposite repertoires of Ebp and AS exhibit divergent biofilm development suggesting that the two adhesins differentially contribute to biofilm structure.

**Figure 4.**
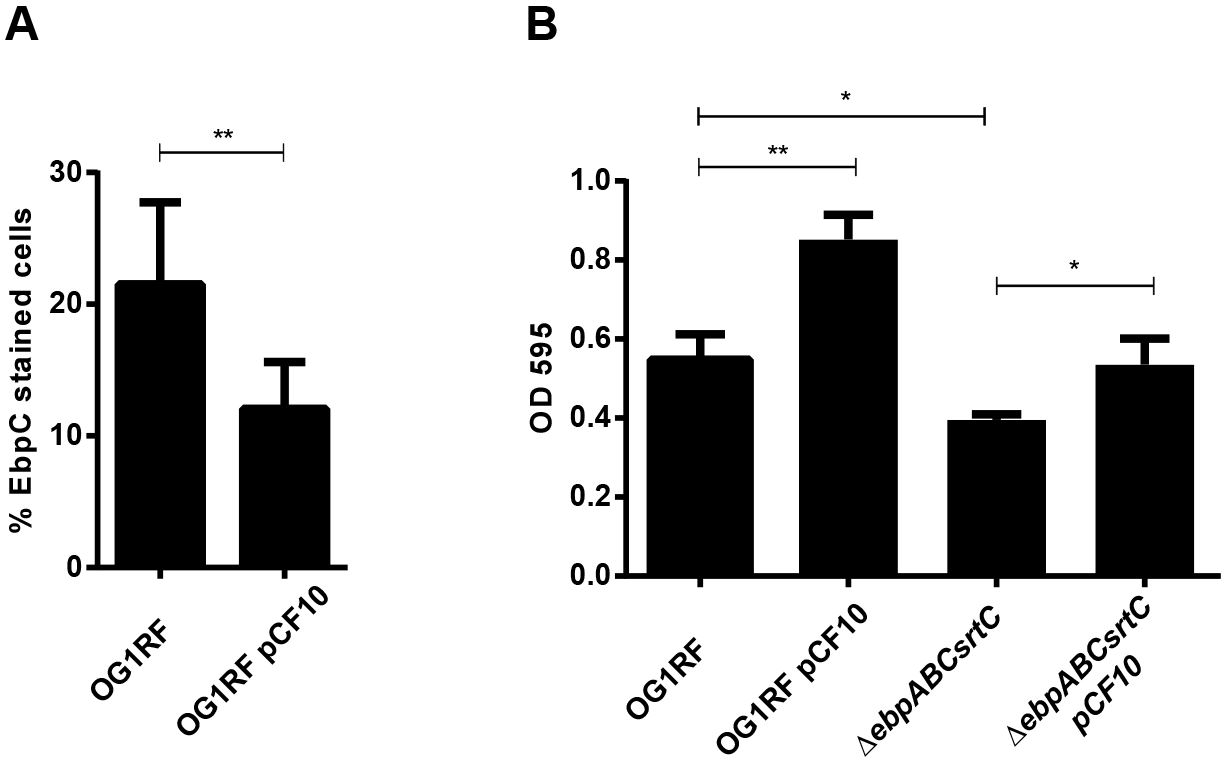
AS contributes to initial attachment and biofilm formation independent of Ebp. **A.** IFM with Ebp antiserum was performed on cells attached to the biofilm chamber 2 hours after induction. Error bars represent the standard deviation from 3 independent experiments. Statistical analysis was performed by unpaired *t-test* using GraphPad * p<0.05, ** p< 0.01, n.s.: p=0.05. **B.** CV staining of biofilms formed by induced (+) and uninduced (−) OG1RF pCF10 and OGlRFΔ*ebpABCsrtC* pCF10 on plastic after 24 hours. Error bars represent the standard deviation from 3 independent experiments. Statistical analysis was performed by unpaired *t-test* using GraphPad * p<0.05, ** p< 0.01.

**Figure 5.**
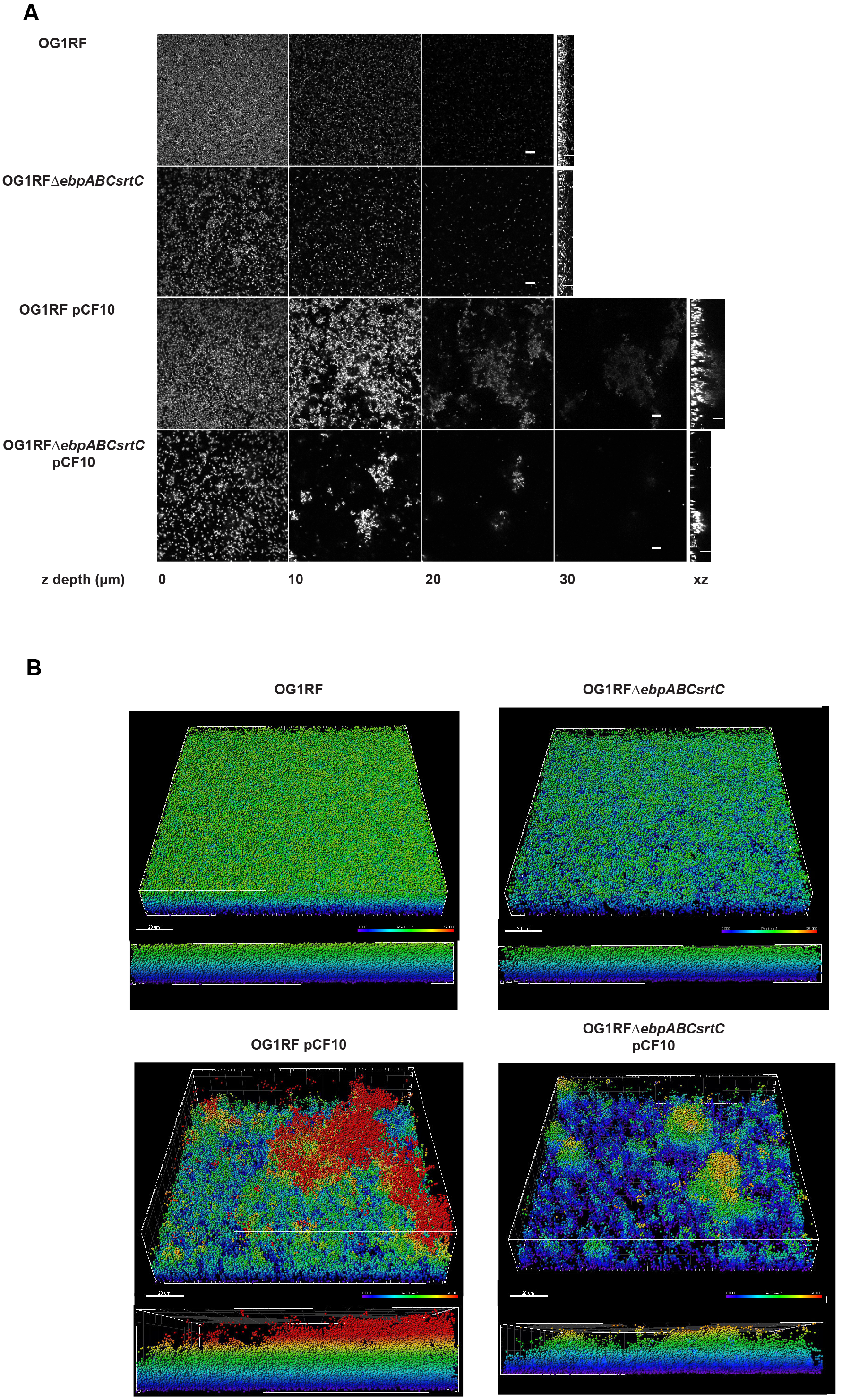
Ebp and AS differentially contribute to biofilm structure and development. Confocal laser scanning microscopy images of 24 hour biofilms grown in plastic chambers in 40% TSBG with cCF10 (1.2 μg/ml). Cells were fixed and stained with Hoechst. **A.** Z stack images represent biofilm depth of 0, 10, 20 and 30 μm, followed by z view for the same samples. Scale bar 10 μm. **B**. Imaris software modelling of the confocal laser scanning microscopy images from **A**, where every cell is represented as a sphere based on Hoechst staining and z-depth is color-coded (0-25 μm) where purple is 0 μm and red is 25 μm. Both perspective (top) and side (bottom) view are shown. Scale bar is 20 μm.

## DISCUSSION

Although most bacteria encode a variety of adhesins for surface attachment and colonization, the number and variety of adhesins expressed in the population and on each cell can vary to maximize adhesive capacities but avoid immune clearance and cross-adhesin interference (23, 30). In *E. faecalis* V583, 17 of 41 predicted Sortase A substrate proteins contain MSCRAMM motifs and at least 7 are expressed within the human host, as antibodies for them are readily detected in the serum (5). This observation raises questions as to when, to what extent, and how promiscuously each adhesin is expressed within the enterococcal population. In the present study, we addressed the relationship between two of these enterococcal adhesins ‒ Ebp and AS.

We showed that Ebp and AS can be co-expressed on the same cell, but that known regulatory mechanisms for each adhesin (EbpR and cCF10, respectively) do not influence expression of the other. However, we found that at low cell density during planktonic growth, simultaneous expression of AS and Ebp interferes with AS-mediated intercellular clumping and reduces HGT rates by 10-fold, presumably due to steric interference by protruding Ebp, which could prevent AS from binding to its receptor, lipoteichoic acid (LTA), on neighboring cells (31). Similarly, *E. coli* type I fimbriae can prevent cellular aggregation mediated by self-associating autotransporters (SAAT), suggesting to the authors that bulky surface proteins such as pili interfere with aggregation and physically mask the function of SAAT-type proteins (32, 33). Therefore pilus-interference with aggregation might be a more general mechanism across bacteria than previously appreciated. Since *E. faecalis* pili are only expressed on a subset of cells, maintaining a subset of unclumped, piliated cells after AS induction may benefit the overall population by limiting the extent to which HGT can occur. Ebp interference may be a general mechanism to protect planktonic cells from abundant plasmid intake which in turn could contribute to genome stability and increased fitness (34). Notably, we can only detect Ebp-dependent differences in HGT rates at early time points, and the difference disappears 2 hours after mating. Similarly, Bhatty et al reported no difference in plasmid transfer rates from WT or Ebp null mutant donors after 2 hours of mating, but they did not examine earlier time points as we did (20). Recently, Ebp on WT donor cells was shown to confer a 1.6-fold increase in conjugative transfer of pAMβ1, after 5 hours of mating, compared to an *ebp* null donor cells (29). While different plasmids, mating times, and experimental cell densities may explain the differing conclusions about the contribution of Ebp to plasmid transfer in these previous studies, coupled with our data, a model emerges in which Ebp may interfere with HGT at low cell densities, but later promote HGT when Ebp-dependent biofilm aggregates start to form which may instead facilitate plasmid transfer. The reasons for these density-dependent differences remain to be determined.

In addition to its impact on HGT, we also demonstrate that co-expression of Ebp and AS may have unique functional contributions to biofilm development. Previously, it was shown that AS acts cooperatively with eDNA to promote biofilm formation resulting in thicker biofilms (20). Here we extend that model, and propose that AS and Ebp differentially contribute to the structure of the biofilm, where AS-mediated clumps of cells that are not expressing Ebp give rise to 3D microcolonies that may be enriched for eDNA. Our data suggests that Ebp-expressing cells can then “fill in” the biofilm space leading to a more densely packed biofilm, as well as contribute the development of thicker 3-D biofilms.

Importantly, we observed similar biomass of OG1RF and Ebp null strain that expresses AS, where AS restores biofilm biomass to OG1RF levels even in the absence of pili. However, despite similarities in biomass accumulation, biofilms formed by only Ebp or AS differed structurally. Taken together our findings suggest that these adhesins may complement each other during biofilm formation and simultaneously contribute different structural roles during biofilm development.

## EXPERIMENTAL PROCEDURES

### Bacterial strains and growth conditions

*Enterococcus faecalis* strains used in this study are listed in **Table 1**. For planktonic growth, strains were grown statically in brain heart infusion media (BHI; BD Difco, USA) or on BHI agar (BHI supplemented with 1.5% agarose (1st BASE, Singapore)) at 37°C. For biofilm assays, tryptone soy broth (Oxoid, UK) supplemented with 10 mM glucose (TSBG) was used. Antibiotics were used in the following concentrations where appropriate: tetracycline (tet), 15 μg/ml; rifampin (rif), 25 μg/ml; streptomycin (str) 500 μg/ml.

**Table 1.** Bacterial strains used in this study.

| Strains | Resistance | Source |
| --- | --- | --- |
| OG1RF | rif, fus, tet | (28) |
| OG1RF pCF10 | rif, fus, tet | This study |
| OG1SS pCF10 | str, tet | (40) |
| OG1XΔ*srtC* | str | (11) |
| OG1RFΔ*ebpABCsrtC* | rif, fus | (9) |
| OG1RFΔ*ebpABCsrtC* pCF10 | rif, fus, tet | This study |
| rif – rifampin, fus – fusidic acid, tet – tetracycline, str - streptomycin |  |  |

### Strain construction

To construct OG1RF pCF10 and OG1RFΔ*ebpABCsrtC* pCF10 we conjugated pCF10 plasmid from OG1SS pCF10 to OG1RF or OG1RFΔ*ebpABCsrtC* as previously described (28) with the following modifications: overnight cultures of OG1SS pCF10 were diluted 1:10 in fresh BHI medium and induced with the peptide pheromone cCF10 (LVTLVFV, 1st Base Peptides, Singapore) at a final concentration of 0.12 ng/mL (35) for 1 hour, with shaking at 200 rpm till optical density at a wavelength of 600 nm (OD_600nm_) 0.3. We then added 0.5 ml of the induced culture to 4.5 ml of mid-log phase culture of statically grown OG1RF or OG1RFΔ*ebpABCsrtC* (overnight cultures were diluted 1:10 in fresh BHI media and grown for approximately 1,5 hours till OD_600nm_ 0.45-0.6). The mixed cultures were incubated at 37°C for 30 minutes, with shaking at 200 rpm before plating on BHI agar containing tet and rif to select for OG1RF pCF10 and OG1RFΔ*ebpABCsrtC* pCF10 transconjugants. Transconjugants were validated by PCR for the presence of the pCF10 plasmid encoded gene *prgB* (both strains) and the absence of *ebpC* (only for OG1RFΔ*ebpABCsrtC*). Primers used are listed in **Table 2**.

**Table 2.**
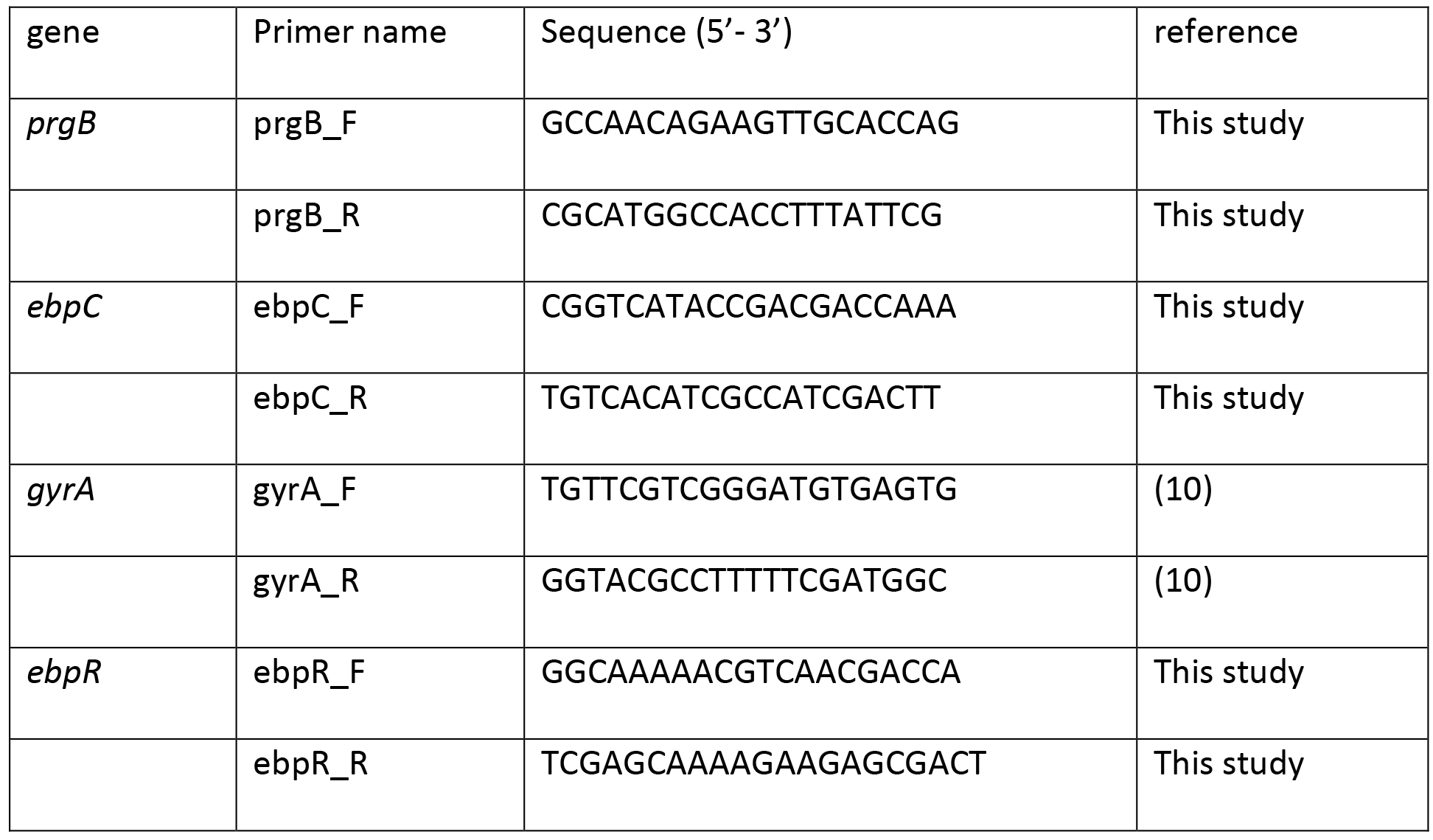
Primers used in this study.

### Horizontal gene transfer (HGT) assay

OG1SS pCF10 donor cells were grown overnight in BHI and then diluted to OD_600nm_ 0.1 in 5 ml of fresh BHI medium in the presence of 0.12 ng/ml of cCF10 peptide. After 2 hours of shaking at 200 rpm at 37°C, visible clumps formed on the bottom of the tube. The top 2 ml represented unclumped (suspension fraction) and 1 ml from the bottom represented the clumped fraction. EDTA at a final concentration of 0.05 M was added to each fraction and vortexed for 15 sec to separate clumped cells (36). Both fractions were normalized to OD_600nm_ 1 and washed with PBS. Donor cells (from the suspension or clumped fraction) were mixed with recipient OG1RFΔ*ebpABCsrtC* cells in a ratio of 1:10 (donor:recipient) and incubated statically at 37°C. After 30 min and 120 min, 1 ml was removed, vortexed for 15 sec, serially diluted, and plated on selective plates to quantify donor OG1SS pCF10 cells on BHI + str + tet and transconjugants OG1RFΔ*ebpABCsrtC* pCF10 on BHI + rif + tet. Alternatively, OG1RF pCF10 and OG1RFΔ*ebpABCsrtC* donor cells were grown as above and were mixed with OG1XAsrtC recipient cells in a ratio of 1:10 (donor:recipient). Conjugation was carried out as above, and cells were plated on selective plates to quantify donor OG1RF pCF10 or OG1RFΔ*ebpABCsrtC* pCF10 cells on BHI + rif + tet agar, and OGlX *AsrtC* pCF10 Tc on BHI + str + tet agar. HGT rates were calculated as the number of Tc per donor cell (Tc/Donor) as described previously (20).

### Generation of polyclonal antisera

Recombinant protein fragments were designed, expressed and purified using Protein Production Platform (NTU, Singapore) with a technology and workflow of the former Biotechnology unit at the Structural Genomics Consortium (Karolinska Institutet, Sweden). Briefly, the AS target (nucleotides 550-1100 from *prgB* NC_006827.2 (12883‥16800)) and Ebp target (nucleotides 1-627 from NC_017316.1 (911134‥913017)) were amplified using multi-construct approach to generate an 184 amino acid and 209 amino acid gene product respectively and were cloned in pNIC28-Bsa4 with an N-terminal His-Tag followed by a TEV protease cleavage site. The resultant plasmid was then transformed to BL21 (DE3) *E. coli* and recombinant protein was expressed following overnight induction with Isopropyl (β-D-1-thiogalactopyranoside. Cells were lysed and recombinant protein was purified by IMAC affinity chromatography using the His-tag followed by a size exclusion chromatography. The purity of the recombinant protein was assessed by SDS-PAGE and mass verified by mass spectrometry. Polyclonal antisera was generated commercially (SABio, Singapore) by immunization of rabbits or guinea pigs with purified recombinant AS or Ebp, respectively. Specificities of the immune sera were confirmed by the absence of signal on Western blots of whole cell lysates from OG1RFΔ*ebpABCsrtC* or pCF10-free strain.

### Immunofluorescent microscopy

To image AS and Ebp on planktonically grown *E. faecalis*, cells were normalized to OD_600_ 1.0, washed once with PBS, and then fixed in 4% paraformaldehyde (PFA) for 20 minutes at 4°C. Cells were again washed, diluted in PBS, and spread on Poly-L-Lysine pre-coated slides (Polysciences Inc, USA). The slides were then blocked with 2% bovine serum albumin (BSA) in PBS and incubated at room temperature (RT) for 20 minutes. After blocking, the slides were washed with PBS and 20 μl of primary antibody (guinea pig anti-Ebp serum, and mouse anti-AS serum) was then added to the slides followed by incubation at 4°C overnight. The next day, the slides were washed 3 times with PBS and 20 μl of fluorescence conjugated secondary antibody (Alexa Fluor 568 goat anti-guinea pig for Ebp and Alexa Fluor 488 goat anti-rabbit for AS) was added and incubated in the dark for 1 hour at RT. Slides were then washed, mounted with VectaShield mounting media (Vector Laboratories Inc, USA) and visualized on an inverted Epi-fluorescence microscope (Zeiss Axio observer Z1, Germany) or Zeiss Elyra PS.1 (Zeiss, Germany) for co-localization studies. The percentage of EbpC- and AS-stained cells was determined by comparing total number of cells viewed under phase contrast to number of stained cells on the merged fluorescent/phase image.

### RNA extraction and RT-qPCR

Bacteria were harvested at the indicated time points by centrifugation at 10,000 rpm for 1 minute. RNA was isolated using the UltraClean^®^ Microbial RNA isolation Kit (Mobio Laboratories Inc, USA) according to manufacturer’s protocol. DNase treatment was performed using the Turbo DNA-free™ kit (Ambion, USA) according to the protocol. RNA concentration and DNA contamination were assessed using a Qubit^®^ 2.0 Fluorometer (Invitrogen, USA). The integrity of the RNA was determined using the TapeStation™ according to manufacturer’s protocol (Agilent Technologies Inc, USA). RNA with RIN values above 7.5 and DNA contamination < 5% was used for RT-qPCR. Equivalent amounts of RNA were converted to cDNA using the SuperScript^®^ III First-Strand Synthesis SuperMix Kit according to the manufacturer’s protocol (Invitrogen, USA). Following cDNA synthesis, RT-qPCR was performed using KAPA SYBR^®^ FAST qPCR Kit MasterMix (2X) Universal according to the manufacturer’s protocol (KAPA Biosystems, USA). The housekeeping gene gyrase A (*gyrA*) was used as an endogenous control in this study as described previously (10, 37). Primers used for amplification of *gyrA*, *ebpC* and *ebpR* are listed in **Table 2**. The ΔΔC_T_ method was used to quantify gene expression differences (38).

### Crystal violet biofilm assay

Crystal violet biofilm assays were performed in a 96 wells plate (Thermofisher, USA) as described previously (39) with the following modifications. Overnight cultures were normalized to OD_600nm_ 0.7, washed and resuspended in 1 ml PBS. 200 μl of the normalized culture was added to 5 ml of TSBG with or without 0.12 ng/ml of cCF10. 200 μl of vortexed cells were seeded into 96-well plates in triplicates and grown at 37°C for 24 hours. Wells were then washed 2 times with PBS and stained with 0.1% crystal violet solution (Sigma-Aldrich, Germany) and incubated for 30 min at 4°C. Wells were then washed three times with PBS and blot dried. 200 μL of ethanol:acetone (80:20) solution was added to each well to solubilize the crystal violet and incubated for 45 min at RT. After incubation, OD_595nm_ readings were taken using a spectrophotometer (UVmini-1240, Shimadzu, Japan).

### Confocal laser scanning microscopy and 3-D modelling

Bacteria were grown and normalized as described above for crystal violet biofilm assay. 200 μl from a suspension of 10^5 cfu/ml in 40% TSBG plus 1.2 ng/ml cCF10 was seeded to 8-well Ibidi chamber slides (Ibidi, Germany) and incubated at 37°C. After 24 hours, supernatants were removed, and remaining cells were fixed with 4% PFA for 20 min at 4°C. The biofilms were then washed once with PBS and DNA was stained with Hoechst 33342 dye (Thermofisher, USA) at a final concentration of 1 μg/ml for 20 min at RT in the dark. We tried both 0.12 and 1.2 ng/ml of pheromone for biofilm assay and while we observed similar clumping for pCF10-carrying strains, the 0.12 biofilms were more delicate and most cells and clumps lifted easily prior PFA staining. Therefore, for consistency in handling we carried out biofilm experiments with 1.2 ng/ml of pheromone. Fixed and stained biofilms were imaged directly from the chambers on a Zeiss LSM780 confocal microscope (Zeiss, Germany) at 60x magnification with a 405 nm laser. Z-stacks were collected through entire biofilm thickness every 0.42 nm before the signal loss. At least 5 z-stacks per chamber were collected at different locations. Fiji software was used to create biofilm montage representations at increments of 5 nm. 3-D reconstruction and modelling was performed with Imaris software (Bitplane, USA), where every cell was represented as 0.5 nm sphere based on DNA staining and z- depth was color-coded.

## ACKNOWLEDGEMENTS

We are grateful to Gary Dunny for providing AS antibody. We thank Pei Yi Choo for assisting with super resolution imaging, and Dr. Artur Matysik and Ling Ning Lam for advice on confocal imaging and data representation. We also thankful to the Protein production platform at the Joint NTU-IMCB center for recombinant proteins production and purification.

## REFERENCES

1. Schlievert PM, Gahr PJ, Assimacopoulos AP, Dinges MM, Stoehr JA, Harmala JW, Hirt H, Dunny GM. 1998. Aggregation and binding substances enhance pathogenicity in rabbit models of Enterococcus faecalis endocarditis. Infection and immunity 66:218–223.

2. Nallapareddy SR, Singh KV, Sillanpaa J, Garsin DA, Hook M, Erlandsen SL, Murray BE. 2006. Endocarditis and biofilm-associated pili of Enterococcus faecalis. J Clin Invest 116:2799–807.

3. Flores-Mireles AL, Pinkner JS, Caparon MG, Hultgren SJ. 2014. EbpA vaccine antibodies block binding of Enterococcus faecalis to fibrinogen to prevent catheter-associated bladder infection in mice. Sci Transl Med 6:254ra127.

4. Navarre WW, Schneewind O. 1994. Proteolytic cleavage and cell wall anchoring at the LPXTG motif of surface proteins in gram-positive bacteria. Mol Microbiol 14:115–21.

5. Sillanpaa J, Xu Y, Nallapareddy SR, Murray BE, Hook M. 2004. A family of putative MSCRAMMs from Enterococcus faecalis. Microbiology 150:2069–78.

6. Olmsted SB, Kao SM, van Putte LJ, Gallo JC, Dunny GM. 1991. Role of the pheromone-inducible surface protein Asc10 in mating aggregate formation and conjugal transfer of the Enterococcus faecalis plasmid pCF10. J Bacteriol 173:7665–72.

7. Toledo-Arana A, Valle J, Solano C, Arrizubieta MaJ, Cucarella C, Lamata M, Amorena B, Leiva J, Penadés JR, Lasa I. 2001. The enterococcal surface protein, Esp, is involved in Enterococcus faecalis biofilm formation. Applied and environmental microbiology 67:4538–4545.

8. Rich RL, Kreikemeyer B, Owens RT, LaBrenz S, Narayana SV, Weinstock GM, Murray BE, Hook M. 1999. Ace is a collagen-binding MSCRAMM from Enterococcus faecalis. J Biol Chem 274:26939–45.

9. Nielsen HV, Flores-Mireles AL, Kau AL, Kline KA, Pinkner JS, Neiers F, Normark S, Henriques-Normark B, Caparon MG, Hultgren SJ. 2013. Pilin and sortase residues critical for endocarditis- and biofilm-associated pilus biogenesis in Enterococcus faecalis. J Bacteriol 195:4484–95.

10. Bourgogne A, Singh KV, Fox KA, Pflughoeft KJ, Murray BE, Garsin DA. 2007. EbpR is important for biofilm formation by activating expression of the endocarditis and biofilm-associated pilus operon (ebpABC) of Enterococcus faecalis OG1RF. J Bacteriol 189:6490–3.

11. Kline KA, Kau AL, Chen SL, Lim A, Pinkner JS, Rosch J, Nallapareddy SR, Murray BE, Henriques-Normark B, Beatty W, Caparon MG, Hultgren SJ. 2009. Mechanism for sortase localization and the role of sortase localization in efficient pilus assembly in Enterococcus faecalis. J Bacteriol 191:3237–47.

12. Bourgogne A, Thomson LC, Murray BE. 2010. Bicarbonate enhances expression of the endocarditis and biofilm associated pilus locus, ebpR-ebpABC, in Enterococcus faecalis. BMC Microbiol 10:17.

13. Nallapareddy SR, Singh KV, Sillanpää J, Zhao M, Murray BE. 2011. Relative contributions of Ebp Pili and the collagen adhesin ace to host extracellular matrix protein adherence and experimental urinary tract infection by Enterococcus faecalis OG1RF. Infection and immunity 79:2901–2910.

14. Nielsen HV, Guiton PS, Kline KA, Port GC, Pinkner JS, Neiers F, Normark S, Henriques-Normark B, Caparon MG, Hultgren SJ. 2012. The metal ion-dependent adhesion site motif of the Enterococcus faecalis EbpA pilin mediates pilus function in catheter-associated urinary tract infection. MBio 3:e00177–12.

15. Waters C, Dunny G. 2001. Analysis of Functional Domains of theEnterococcus faecalis Pheromone-Induced Surface Protein Aggregation Substance. Journal of bacteriology 183:5659–5667.

16. Nakayama J, Ruhfel RE, Dunny GM, Isogai A, Suzuki A. 1994. The prgQ gene of the Enterococcus faecalis tetracycline resistance plasmid pCF10 encodes a peptide inhibitor, iCF10. J Bacteriol 176:7405–8.

17. Chandler JR, Hirt H, Dunny GM. 2005. A paracrine peptide sex pheromone also acts as an autocrine signal to induce plasmid transfer and virulence factor expression in vivo. Proceedings of the National Academy of Sciences 102:15617–15622.

18. Olmsted S, Erlandsen S, Dunny G, Wells C. 1993. High-resolution visualization by field emission scanning electron microscopy of Enterococcus faecalis surface proteins encoded by the pheromone-inducible conjugative plasmid pCF10. Journal of bacteriology 175:6229–6237.

19. Chuang-Smith ON, Wells CL, Henry-Stanley MJ, Dunny GM. 2010. Acceleration of Enterococcus faecalis biofilm formation by aggregation substance expression in an ex vivo model of cardiac valve colonization. PLoS One 5:e15798.

20. Bhatty M, Cruz MR, Frank KL, Gomez JA, Andrade F, Garsin DA, Dunny GM, Kaplan HB, Christie PJ. 2015. Enterococcus faecalis pCF10-encoded surface proteins PrgA, PrgB (aggregation substance) and PrgC contribute to plasmid transfer, biofilm formation and virulence. Mol Microbiol 95:660–77.

21. Hirt H, Schlievert PM, Dunny GM. 2002. In vivo induction of virulence and antibiotic resistance transfer in Enterococcus faecalis mediated by the sex pheromone-sensing system of pCF10. Infection and immunity 70:716–723.

22. Chuang ON, Schlievert PM, Wells CL, Manias DA, Tripp TJ, Dunny GM. 2009. Multiple functional domains of Enterococcus faecalis aggregation substance Asc10 contribute to endocarditis virulence. Infection and immunity 77:539–548.

23. Xia Y, Gally D, Forsman-Semb K, Uhlin BE. 2000. Regulatory cross-talk between adhesin operons in Escherichia coli: inhibition of type 1 fimbriae expression by the PapB protein. The EMBO Journal 19:1450–1457.

24. Kazmierczak BI, Schniederberend M, Jain R. 2015. Cross-regulation of Pseudomonas motility systems: the intimate relationship between flagella, pili and virulence. Curr Opin Microbiol 28:78–82.

25. Holden NJ, Totsika M, Mahler E, Roe AJ, Catherwood K, Lindner K, Dobrindt U, Gally DL. 2006. Demonstration of regulatory cross-talk between P fimbriae and type 1 fimbriae in uropathogenic Escherichia coli. Microbiology 152:1143–53.

26. Olmstead S, Erlandsen S, Dunny G, Wells C. 1993. High-resolution visualization by field emission scanning electron microscopy of Enterococcus faecalis cell surface proteins encoded by the pheromone-induced conjugative plasmid pCF10. J Bacteriol 175:6229–6237.

27. Chatterjee A, Cook LC, Shu CC, Chen Y, Manias DA, Ramkrishna D, Dunny GM, Hu WS. 2013. Antagonistic self-sensing and mate-sensing signaling controls antibiotic-resistance transfer. Proc Natl Acad Sci U S A 110:7086–90.

28. Dunny GM, Brown BL, Clewell DB. 1978. Induced cell aggregation and mating in Streptococcus faecalis: evidence for a bacterial sex pheromone. Proceedings of the National Academy of Sciences 75:3479–3483.

29. La Rosa SL, Montealegre MC, Singh KV, Murray BE. 2016. Enterococcus faecalis Ebp Pili are Important for Cell-Cell Aggregation and Intraspecies Gene Transfer. Microbiology doi:10.1099/mic.0.000276.

30. Kline KA, Falker S, Dahlberg S, Normark S, Henriques-Normark B. 2009. Bacterial adhesins in host-microbe interactions. Cell Host Microbe 5:580–92.

31. Waters CM, Hirt H, McCormick JK, Schlievert PM, Wells CL, Dunny GM. 2004. An amino-terminal domain of Enterococcus faecalis aggregation substance is required for aggregation, bacterial internalization by epithelial cells and binding to lipoteichoic acid. Mol Microbiol 52:1159–71.

32. Klemm P, Vejborg RM, Sherlock O. 2006. Self-associating autotransporters, SAATs: Functional and structural similarities. International Journal of Medical Microbiology 296:187–195.

33. Sherlock O, Schembri MA, Reisner A, Klemm P. 2004. Novel Roles for the AIDA Adhesin from Diarrheagenic Escherichia coli: Cell Aggregation and Biofilm Formation. Journal of Bacteriology 186:8058–8065.

34. Starikova I, Al-Haroni M, Werner G, Roberts AP, Sørum V, Nielsen KM, Johnsen PJ. 2013. Fitness costs of various mobile genetic elements in Enterococcus faecium and Enterococcus faecalis. Journal of Antimicrobial Chemotherapy 68:2755–2765.

35. Antiporta MH, Dunny GM. 2002. ccfA, the Genetic Determinant for the cCF10 Peptide Pheromone in Enterococcus faecalis OG1RF. Journal of Bacteriology 184:1155–1162.

36. Chuang-Smith ON, Wells CL, Henry-Stanley MJ, Dunny GM. 2010. Acceleration of Enterococcus faecalis biofilm formation by aggregation substance expression in an ex vivo model of cardiac valve colonization. PLoS One 5:e15798.

37. Lebreton F, Riboulet-Bisson E, Serror P, Sanguinetti M, Posteraro B, Torelli R, Hartke A, Auffray Y, Giard J-C. 2009. ace, which encodes an adhesin in Enterococcus faecalis, is regulated by Ers and is involved in virulence. Infection and immunity 77:2832–2839.

38. Livak KJ, Schmittgen TD. 2001. Analysis of relative gene expression data using real-time quantitative PCR and the 2(-Delta Delta C(T)) Method. Methods 25:402–8.

39. Keogh D, Lam LN, Doyle LE, Matysik A, Pavagadhi S, Umashankar S, Low PM, Dale JL, Song Y, Ng SP. 2018. Extracellular Electron Transfer Powers Enterococcus faecalis Biofilm Metabolism. mBio 9:e00626–17.

40 Franke AE, Clewell DB. 1981. Evidence for a chromosome-borne resistance transposon (Tn916) in Streptococcus faecalis that is capable of “conjugal” transfer in the absence of a conjugative plasmid. J Bacteriol 145:494–502.

